# revtools: An R package to support article screening for evidence synthesis

**DOI:** 10.1101/262881

**Authors:** Martin J. Westgate

## Abstract

The field of evidence synthesis is growing rapidly, with a corresponding increase in the number of software tools and workflows to support the construction of systematic reviews, systematic maps, and meta-analyses. Despite much progress, however, a number of problems remain including slow integration of new statistical or methodological approaches into user-friendly software, low prevalence of open-source software, and poor integration among distinct software tools. These issues hinder the utility and transparency of new methods to the research community. Here I present revtools, an R package to support article screening during evidence synthesis projects. It provides tools for the import and de-duplication of bibliographic data, screening of articles by title or abstract, and visualization of article content using topic models. The software is entirely open-source and combines command-line scripting for experienced programmers with custom-built user interfaces for casual users, with further methods to support article screening to be added over time. Revtools provides free access to novel methods in an open-source environment, and represents a valuable step in expanding the capacity of R to support evidence synthesis projects.

## Introduction

Evidence synthesis is increasingly used to help inform policy responses to critical societal issues ^1^. Further, the growth of evidence synthesis methodologies – predominantly systematic reviews (SRs), but also including systematic maps and meta-analyses – has been concurrent with a massive increase in the amount of published scientific material, estimated at 50 million articles in 2010 ^2^ and now growing at over 2.5 million articles a year ^3^. In combination, these dual pressures have focused attention on new methods to support SR, with a vast number of software tools and workflows available for researchers ^4, 5^. Despite much progress, however, several problems have emerged in the SR software landscape during this rapid expansion. In particular, many software tools have limited interoperability with one another, or hide the details of the algorithms they provide to assist with article screening and related tasks, problems that are incentivized in commercial environments. Further, new statistical or computational methods are not always made available to users in a timely or easily-accessible way, hindering their application to real problems ^6^. Addressing these issues would greatly assist progress in the application of novel techniques to evidence synthesis projects ^7^.

In recent years, an alternative software development model has emerged which draws on community-maintained programming languages such as R or Python to support bespoke, open-source software tools. These tools have the advantage that they can be provided free, and often more quickly than they can be integrated into commercial software ^8^. Further, the open source nature of these tools means they are able to build on vast numbers of software libraries built by other members of the development community, allowing rapid adaptation and specialization. While these changes have had some impact on evidence synthesis workflows – and particularly on meta-analysis, where many software tools are now available ^9^ – there has been more limited development of tools to support data search and screening. This is problematic because those tasks form the foundation of effective systematic reviews.

In this paper I present *revtools*, an R package for importing, manipulating and screening bibliographic data. The package serves a dual function of providing robust infrastructure to support current systematic reviews and meta-analyses, while also supporting the development of more advanced software tools, including novel data visualization tools to facilitate article screening (see Table 1 for a description of key functions). Although developed entirely within a command-line-based environment, *revtools* has been designed to be as user-friendly as possible by including interfaces that launch in the users’ web browser. These interfaces are built using the R packages *Shiny* ^10^ and *shinydashboard* ^11^, removing any dependencies on external software. Functions are also provided to help experienced programmers to explore the underlying models and data in more detail. Users can download the stable version of *revtools* from CRAN (https://cran.r-project.org/package=revtools); view, download or contribute to the development version on GitHub (https://github.com/mjwestgate/revtools); or view further documentation and examples on a dedicated website (https://revtools.net). Below I describe the functionality provided by *revtools* in order of a typical SR workflow.

**Table 1:** Key functions available in *revtools*

| Category | Function | Description |
| --- | --- | --- |
| Data manipulation | *read_bibliography* | Import bibliographic data from a range of formats into an R *data.frame* |
|  | *write_bibliography* | Export bibliographic data from R into *.ris* or *.bib* formats |
|  | *merge_columns* | Combine (*rbind*) two *data.frames* with different columns into a single *data.frame* |
| Duplicate detection | *find_duplicates* | Identify potentially duplicated references within a *data.frame* |
|  | *extract_unique_references* | Return a *data.frame* that contains only ‘unique’ references |
|  | *screen_duplicates* | Manually screen potential duplicates identified via *find_duplicates* |
| Text mining and visualisation | *make_dtm* | Take a vector of strings and return a Document-Term Matrix (DTM) |
|  | *run_topic_model* | Run a topic model with specified properties |
|  | *screen_topics* | Use topic models to visualize and screen articles |
| Distributing tasks among a team | *allocate_effort* | Assign proportions of articles to members of a team |
|  | *distribute_tasks* | Use results from *allocate_effort* to split a *data.frame*, returning a *.csv* file for each reviewer |
|  | *aggregate_tasks* | Combined screened references back into a single *data.frame* |
| Manual screening | *screen_titles* | Screen articles by title |
|  | *screen_abstracts* | Screen articles by abstract |

## Functionality

### Data import and export

When conducting a systematic review, it is common to have bibliographic data stored in files generated by a bulk download of some kind, typically from an academic database such as Web of Science or Scopus. Bibliographic data from these sources can be stored in a range of formats (with BibTeX and RIS formats among the most common), and while there are R packages for importing BibTeX data (e.g. RefManageR ^12^), there is no function to identify the format of a bibliographic data file and import it in a consistent manner. This is problematic for tagged styles for which no standard formatting exists, for example where two files have identical content but different filename suffixes (such as .*ciw* and .*ris*), or have the same suffix but different formats (common with .*ris*, for example). In *revtools*, the function *read_bibliography* auto-detects tag data and article delimiters, and converts them into a *data*.*frame* (the standard format for R datasets) with human-readable column names. If the dataset is already stored in a spreadsheet-like *csv* file, then *read_bibliography* imports the file as is, with a small number of changes to ensure consistency with other formats. Finally, users can export bibliographic data from R using the function *write_bibliography*.

Although *read_bibliography* returns a *data*.*frame* by default, it does so by means of an intermediate object stored in the new class *bibliography*. Accessing this data class is optional and will not be necessary for most users, but is included because its list-based structure allows easy storage of nested information structures, as is common in bibliographic datasets. Specifically, each *bibliography* object is a list in which each entry is an article, consisting of another list storing standard bibliographic data such as the title, author or keywords for that reference. A priority for future development is to facilitate deeper levels of nestedness where required, such as data on author affiliations or ORCID ID numbers. That functionality would be very useful for integrating bibliographic datasets with (co)citation data, in the context of ‘research weaving’ projects, for example ^13^. In anticipation of this added functionality, class *bibliography* is fully supported as an S3 class with its own *print, summary*, and subset (‘[’ and ‘[[’) functions, and can be converted to a *data*.*frame* using *as*.*data*.*frame* or back again using *as*.*bibliography*.

### Identification and removal of duplicates

A key component of systematic review protocols is that researchers should search a number of databases to ensure that all available literature is detected ^14^. This approach is very thorough, but has the side effect of introducing duplicates into the dataset that must be removed prior to article screening. Rather than building an automated function that makes many decisions on behalf of the user (which is difficult to optimize ^15^), *revtools* implements a fuzzy-matching algorithm for duplicate detection. This approach is more general and flexible but requires some care to return sensible results.

The main function for identifying duplicates in *revtools* is *screen_duplicates*, which runs a GUI in the browser that allows calculation and manual screening of duplicates. Alternatively, the user can call the underlying function – called *find_duplicates* – directly from the workspace, as both functions use the same arguments. First, the user selects a variable from their dataset that they want to investigate for potential duplicates, such as the article titles or DOIs. Second, they select any ‘grouping’ variables that defines a subset of the data where duplicates should be sought. For example, the user may wish to only seek matching titles within the same journal, or the same publication year. Leaving this argument blank leads to more duplicates being located but takes longer to run. Third, the user specifies a method for matching potential duplicates; either an exact match, or fuzzy matching using either *stringdist* ^16^ or the new function *fuzzdist*, which implements functions from the *fuzzywuzzy* Python library (https://github.com/seatgeek/fuzzywuzzy). Finally, the user can set the threshold for deciding whether two strings (which may contain titles, DOIs etc.) are identical, and choose whether to convert strings to lower case or to remove punctuation. If the user has called *screen_duplicates*, setting these arguments and hitting the ‘calculate duplicates’ button will lead to a display of potential duplicated entries, allowing the user to interactively choose between them. If instead they have called *find_duplicates*, the function will return a vector where duplicated articles share the same value. The user can then pass this vector to *screen_duplicates* for manual screening, or use *extract_unique_references* to attempt the de-duplication for them. Future versions of *revtools* will supplement these methods with further options for automation of article de-duplication.

### Visualizing search results using text mining

A common problem in systematic reviews is that search results from academic databases nearly always contain information that is not relevant to the target question. While tools to improve the sensitivity and specificity of article searches exist, this problem is difficult to avoid entirely because many fields lack well-defined tagging ontologies to improve search efficiency (with Medical Subject Headings [MeSH] being the obvious exception), while semantic properties of natural languages ensure that any given keyword is likely to be used in a range of contexts ^17^. What is needed, therefore, is a method which allows the user to gain an *a priori* impression of the dominant themes within their search results ^18^, and to filter out irrelevant content prior to manual screening.

In *revtools*, the new function *screen_topics* attempts to address this gap by visualizing the results of a topic model ^19^ run on user-selected bibliographic data. This allows the user to decide how many themes (or ‘topics’) they would like to identify within the corpus, and which data should be used to parameterize the model (typically some combination of titles, keywords or abstracts). Once the model has been calculated, the main window shows a scatterplot in which each point represents one article, with points colored according to the highest-weighted topic for that article (Fig. 2). Also shown is a bar chart showing the number of articles within each topic, which also serves as a key to the words that make up those topics. If article abstracts are available in the dataset, these are shown below the main panel. Finally, an expandable sidebar gives access to user options, including changing between article or word displays, changes to the color scheme using the viridis family of palettes ^20^, or saving progress in a range of formats. By default, the *screen_topics* uses articles as its unit of analysis; but the user can change this behavior, for example to look for themes among journals or years.

The main value of *screen_topics* is in interacting with bibliographic data in an intuitive way to exclude irrelevant material, using diagrams drawn using the *plotly* R package ^21^. Points or bars that are selected bring up further information that can be used to select or exclude a given article, word, or topic. One critical feature is that all data are linked, meaning that the user can exclude uninformative content and re-run the topic model at any time, unlike most related tools that require topic models to be pre-calculated (e.g. *LDAvis* ^22^). While useful, this approach can lead to slow performance when models are recalculated, even accounting for default settings that prioritize speed over accuracy (i.e. 5 topics, 10,000 iterations). These settings are adequate for a cursory investigation, but users that wish to exclude groups of articles based on their assigned topic, or that require robust classification of the whole corpus, will need to increase these values. More broadly, using topics as a basis for article inclusion or exclusion is subjective and may introduce bias. This approach should only be used for classifying articles in a very broad sense – such as whether they are within or outside of the target discipline – and should not be considered a replacement for careful screening of individual articles during systematic review.

If users wish to explore topic models in more detail, they can run their own models using *run_topic_model*. This is currently a wrapper function to the *topicmodels* library ^23^, but other methods of topic identification will be added in future. The main input to *run_topic_model* is a document-term matrix, which can be constructed using the new function *make_dtm*. This function takes the user-specified text data and applies several standard text processing functions from the *tm* package ^24^, namely: conversion to lower case; removal of punctuation, numbers and ‘stop’ words (such as ‘the’, ‘and’ etc.); stemming; and the removal of short words (<3 letters) or rare terms (present in <1% of articles). After collating all words that contain the same stem, this function returns whichever of the original terms was most common in the dataset, rather than the stem itself (i.e. ‘manag’ might be returned as ‘management’). These functions allow users to duplicate the methods used by *screen_topics* in the R workspace and facilitate increased flexibility to explore model performance, for example by comparing the fit of different model types, or models containing different numbers of topics. This is useful if the topic model and its results are themselves the focus of the investigation ^25^.

### Dividing screening tasks among a team

Systematic reviews are often conducted by teams of researchers who work together to ensure high standards within the screening process. A common approach is ‘double-screening’; allocating each article to two separate reviewers and comparing the results, which helps to ensure consistency in the application of screening criteria and that disagreements about article relevance are discussed openly, leading to reduced bias ^14^. In large reviews, however, or where time is limited, teams may instead choose to use partial double-screening, whereby only a subset of articles are double-screened and the remainder screened once only. Either way, it is important that teams of reviewers have a means of dividing a dataset of articles among participating researchers.

In *revtools*, users can decide how much work they will each perform using the function *allocate_effort*, including specifying the size of their team, the proportion of articles to be checked by more than one reviewer, and the maximum number of times each article should be checked. It is also possible – though optional – to specify reviewer names and to allocate different proportions of articles to each reviewer. Users can either call *allocate_effort* directly, or via the higher-level function *distribute_tasks*, which also requires a bibliographic dataset as input and (by default) outputs a set of .*csv* files with the required properties. Although the user sets the proportion of articles to be reviewed by each person, the identity of the articles in each subset is chosen at random, and so is sensitive to external settings such as *set*.*seed*. Once the reviews have been completed (using *screen_abstracts*, for example), they can be re-combined into a single *data*.*frame* using the *aggregate_tasks* function.

### Manual article screening

The final stage of the systematic review process that can be completed using *revtools* is manual screening of articles to determine their relevance to the review. Users can choose to screen by title only using *screen_titles*, or to screen article titles and abstracts simultaneously using *screen_abstracts*. The latter function is superficially similar to related tools in the *metagear* package ^26^, with the exception that the *revtools* equivalent is presented using *Shiny* for consistency with *screen_duplicates* and *screen_topics*, and to reduce dependence on external software. Both interfaces have broadly similar content: they both allow the user to drag-and-drop a dataset into the app for screening; the user can choose to show or hide information that identifies the author and journal for consistency with SR guidelines; and they can choose the order in which articles are presented. The text colour changes to reflect the users’ decisions as articles are selected or excluded, and both apps allow the user to export their decisions to an external file, or back to the workspace for further processing in R. The only major difference between them is in their displays; *screen_abstracts* shows only one article at a time, whereas by default *screen_titles* shows eight titles at once (Fig. 1).

**Figure 1:**
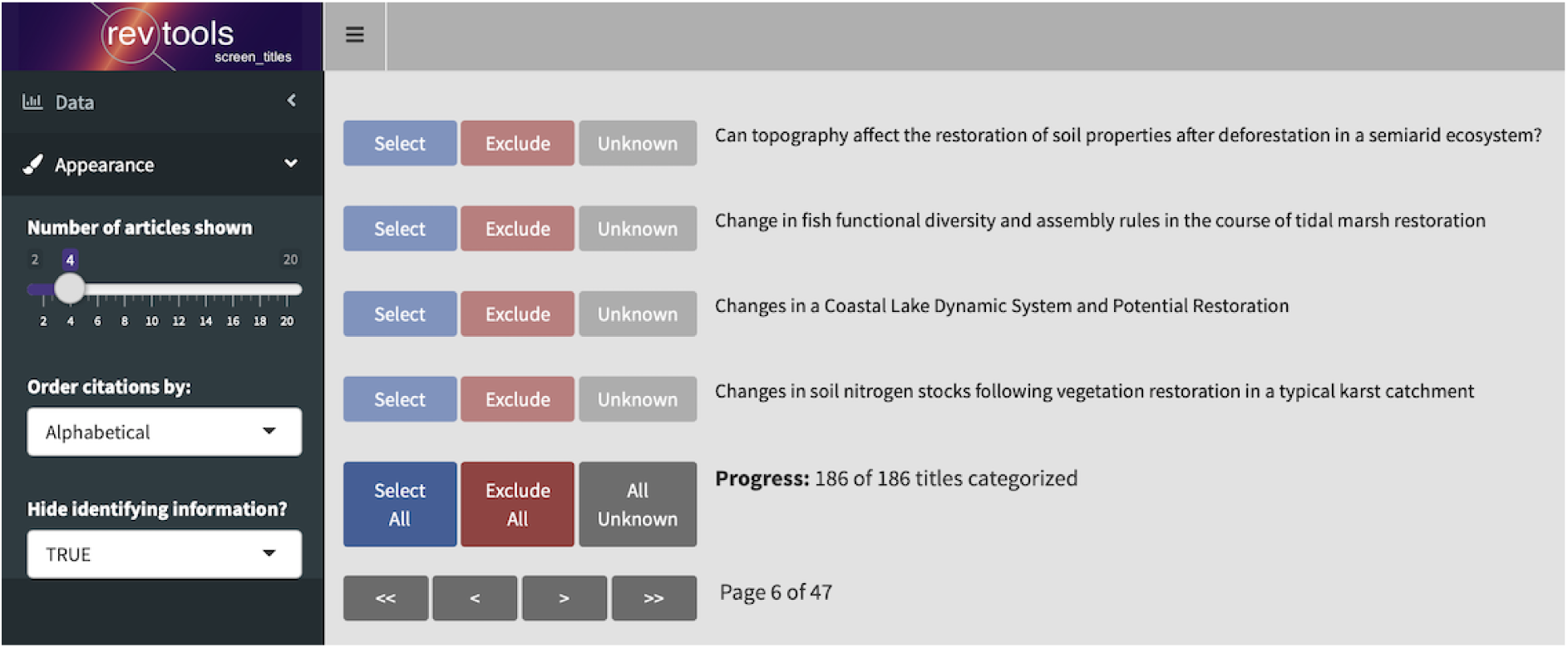
Screenshot of the *screen_titles* interface. Article titles are taken from the example ‘restoration’ dataset (see text).

## Example: Screening diverse datasets

To demonstrate the capabilities available in *revtools*, here I provide a worked example of how the tools described above may be integrated to solve real problems in evidence synthesis projects. To generate an example dataset, I searched Web of Science, Scopus and PubMed on 30^th^ January 2019 for all articles containing the term ‘restoration’ in the title that had been published so far that year. These searches returned 247, 286 and 104 articles respectively, or 637 articles in total. The term ‘restoration’ was chosen as it was given as an example of homonymy in a related paper on article classification ^27^, while the time limits were chosen as a simple means of reducing the size of the dataset for demonstration purposes. A real systematic review search would include a more sophisticated combination of search terms and would not be restricted to such a narrow time window.

The analysis begins by installing revtools, and then reading the three datasets into R using read_bibliography, as follows:

~~~
# install and load package
install.packages (“revtools”)
library(revtools)
# import data from working directory
data_list <- read_bibliography(list.files())
# retain only columns that with >300 non-missing values
keep_cols <- which(
  apply(data_unique, 2, function(a){
    length(which(!is.na(a)))
  }) > 300
)
data <- data[, keep_cols]
~~~

As there is a high likelihood that these three databases will contain some overlapping references, the next logical step is to search for and remove those duplicates. We could achieve this interactively using *screen_duplicates*, or in the command line as follows:

~~~
# find duplicated DOIs within the dataset
doi_match <- find_duplicates(data,
match_variable = “doi”,
group_variables = NULL,
match_function = “exact”
)
# automatically extract one row per duplicate
data_unique <- extract_unique_references(data, doi_match)
nrow(data_unique) # n = 491, i.e. duplication rate = 23%
# an alternative is to try fuzzy title matching
title_match <- find_duplicates(data,
  match_variable = “title”,
  group_variables = NULL,
  match_function = “stringdist”,
  method = “osa”,
  threshold = 5
)
length(unique(title_match)) # n = 492; one less than DOI
# but also slower than exact matching, especially for large datasets
~~~

The final stage is to use *screen_topics* to look for themes in the corpus. Importantly, if we specify an object for this function to return to, then we can save and reload our progress from within R, as follows:

~~~
result <-screen_topics(data_unique)
~~~

In this case, we conduct a journal-level analysis, and include article titles, abstracts and keywords in our topic model. Figure 2 shows the resulting ordination after calculation 10 topics and removing a set of additional stopwords, as well as the original search term (‘restoration’).

**Figure 2:**
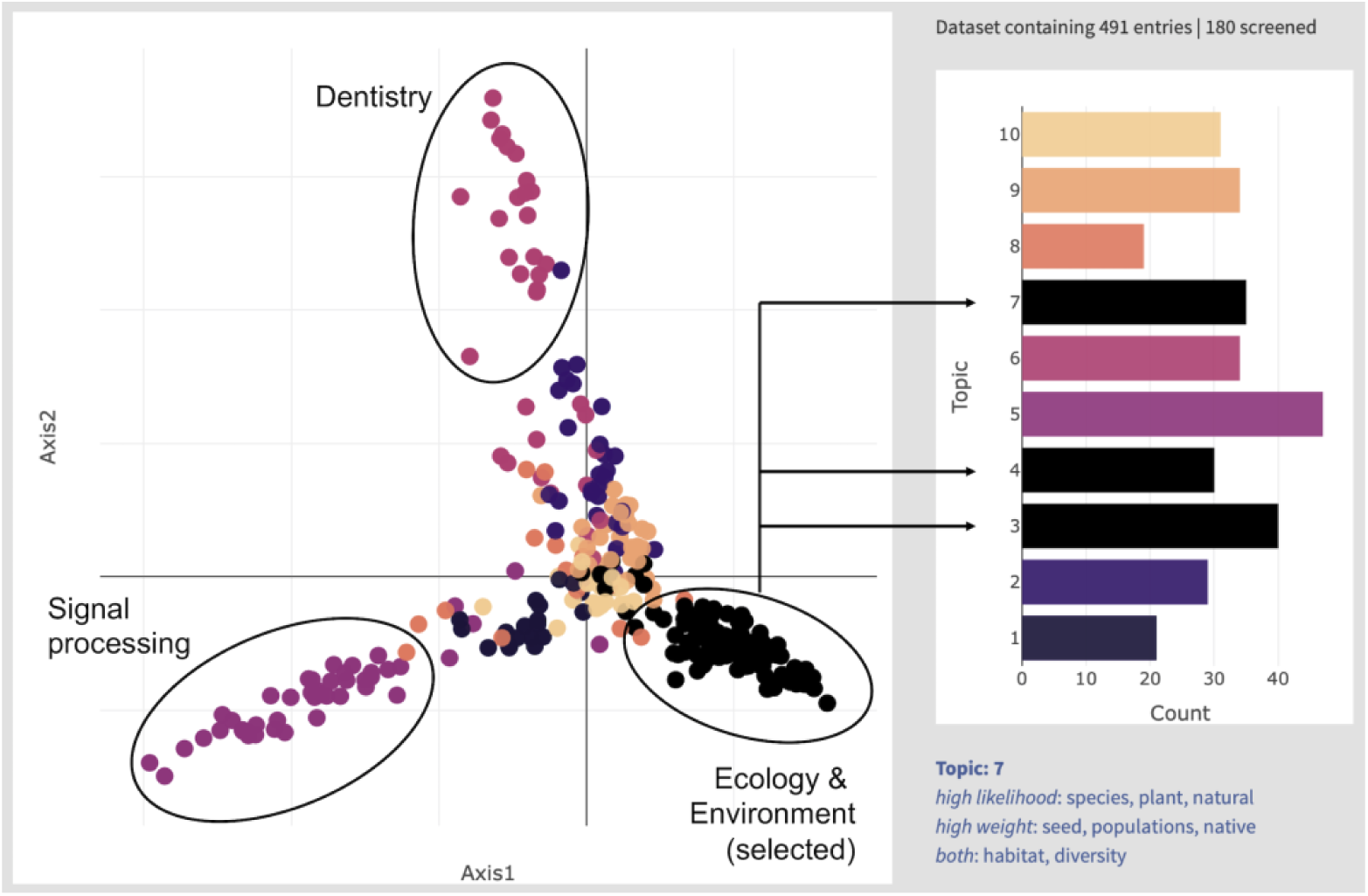
Annotated screenshot of *screen_topics* on the ‘restoration’ dataset, showing one point per journal and ten topics. Topics containing environmental keywords have been selected for further screening and are shown in black.

This analysis shows that a number of discrete disciplines are represented – including engineering, medicine, signal processing and dentistry – while three separate topics in this ordination refer to different uses of the term ‘restoration’ in ecological or environmental contexts. Using the ‘topics’ barplot in *screen_topics* (right-hand side of Fig. 2) we can ‘select’ these topics as being of further interest, while leaving the remaining topics uncategorized. Given the high degree of overlap among topics in the centre of the scatterplot, however, a more risk-averse alternative might be to exclude clearly irrelevant topics (such as signal processing and dentistry, topics 5 & 6 respectively) and manually screen the remaining articles. By using the ‘exit app’ button in the ‘data’ tab of the sidebar, these selections are passed back to the workspace. The resulting object (of class ‘*screen_topics_progress*’) contains our selections as an additional column added on to our source dataset, and are stored in the section named ‘raw’:

~~~
articles_data <-result$raw
env_articles <-articles_data[which(articles_data$selected),]
nrow(env_articles) # n = 186
~~~

It is then a simple process to screen these articles manually using *screen_titles* or *screen_abstracts*, or to investigate the themes within this subset of articles in more detail using *screen_topics* for a second time.

## Discussion

In this paper I have presented *revtools*, a new R package to support article screening for evidence synthesis projects. As well as providing basic infrastructure such as support for a range of bibliographic data formats and tools for de-duplication and article screening, *revtools* supports tools that draw on text mining to explore patterns in bibliographic data. These tools can rapidly summarize topics and locate relevant information within any set of article abstracts, irrespective of academic discipline.

While tools for visualizing text data are not yet common in systematic reviews, expanding their use will be critical if systematic reviews are to remain viable given projected increases in the size of the academic literature. Research from the environmental sciences shows that the average systematic review (SR) currently requires 164 person-days of work ^28^. While comparable data are not available for medical SRs, current data suggest that these take an average of five people 67 weeks to complete ^29^. Further increases the number of articles would make these approaches unviable for most researchers or teams; indeed some authors have argued that this has already happened, specifically for understanding impacts of rare interventions for biodiversity conservation ^30^. Without access to robust time-saving software tools, evidence synthesis researchers risk facing a marked increase in the so-called ‘synthesis gap’; an incapacity of the scientific establishment to synthesize its’ own research for application to societal problems ^7^.

While *revtools* is already a well-developed software package, there are a number of ways in which its functionality could be expanded to improve efficiencies in evidence synthesis. One priority is to allow users to test the effect of fitting covariates to topic models as implemented in the *stm* R package ^31^; another is to explore different methods of displaying articles in 2D space, such as the t-SNE algorithm ^32^. Both of these approaches could address one shortcoming of the current topic model infrastructure in *revtools*, namely that these models take a long time to run, particularly for large datasets. A more advanced feature would be to implement some form of dynamic article recommendation, whereby user choices are used to inform recommendations about potentially relevant information. This approach is widely accepted ^33^ and a range of implementations exist ^27, 34^, although few of them have been rigorously tested ^6^. Besides making these tools more widely available, therefore, one advantage of incorporating them into R would be to facilitate comparisons amongst these distinct but related approaches. Finally, access to the tools provided by *revtools* would be greatly improved if they were available outside of R, a feature made more achievable by their construction as *Shiny* apps that are relatively easy to host online. This development would greatly increase access by users unwilling or unable to work in a scripting environment, and will be a priority for future software releases.

Despite plans to expand the functionality of *revtools*, it is not intended that it should become a standalone package capable of supporting the entire evidence synthesis pathway. Instead, the features provided by *revtools* are likely to be particularly powerful when used in combination with those of related packages. Currently, for example, it is possible to plan a search strategy with *litsearchr* ^*35*^; screen your articles with *revtools*; download articles using *fulltext* ^36^; analyze the resulting data in *metafor* ^37^ or one of the many other meta-analysis packages available in R ^9^; and present a database of study results using *EviAtlas* ^38^. While this is encouraging, many stages of the systematic review workflow remain unsupported within R, with data extraction being a key gap. To address this, several of the developers of the packages listed above are working together to construct an integrated package named *metaverse* to guide users through a standard systematic review workflow. This project will also focus on building new software to integrate existing tools and fill the gaps between them. It is intended, therefore, that *revtools* will continue to grow, but retain a focus on article screening and visualization of textual data sources.

## Acknowledgements

N. Haddaway contributed substantial discussion and insight regarding the design of *revtools*. Those discussions were supported by an International Outgoing Visiting Fellowship to MW, awarded by the Australian Research Councils’ Centre of Excellence for Environmental Decisions. Comments from P. Barton and three anonymous reviewers greatly improved this manuscript.

